# A comparative analysis of Chikungunya and Zika transmission

**DOI:** 10.1101/078923

**Authors:** Julien Riou, Chiara Poletto, Pierre-Yves Boëlle

## Abstract

The recent global dissemination of Chikungunya and Zika has fostered public health concern worldwide. To better understand the drivers of transmission of these two arboviral diseases, we propose a joint analysis of Chikungunya and Zika epidemics in the same territories, taking into account the common epidemiological features of the epidemics: transmitted by the same vector, in the same environments, and observed by the same surveillance systems. We analyse eighteen outbreaks in French Polynesia and the French West Indies using a hierarchical time-dependent SIR model accounting for the effect of virus, location and weather on transmission, and based on a disease specific serial interval. We show that Chikungunya and Zika have similar transmission potential in the same territories (transmissibility ratio between Zika and Chikungunya of 1.04 [95% credible interval: 0.97; 1.13]), but that detection and reporting rates were different (around 19% for Zika and 40% for Chikungunya). Temperature variations between 22°C and 29°C did not alter transmission, but increased precipitation showed a dual effect, first reducing transmission after a two-week delay, then increasing it around five weeks later. The present study provides valuable information for risk assessment and introduces a modelling framework for the comparative analysis of arboviral infections that can be extended to other viruses and territories.

## 1. Introduction

Arboviral infections are increasingly becoming a global health problem [1]. Dengue fever and yellow fever viruses have been re-emerging in many tropical areas since the 1980s [2], but new epidemic waves have recently been caused by lesser known arboviruses: the Chikungunya virus (CHIKV) since 2005 [3], and the Zika virus (ZIKV) since 2007 [4]. Interestingly, the spread of ZIKV and CHIKV have shared many epidemiological characteristics. While discovered in the 1940-50s, the global spread of these viruses to previously unaffected areas has only begun in recent years, and large outbreaks have affected the immunologically naive populations of the Indian and Pacific oceans and of the Americas [5, 6, 7]. Case identification and counting has been an issue for epidemiological surveillance since symptoms caused by ZIKV and CHIKV infection are most of the times mild and not specific. Finally, both diseases can be transmitted by the same mosquitoes of the *Aedes* genus [8, 9]. The most common vector, *Ae. aegypti*, is well adapted to the human habitat [10], is resistant to many insecticides [11], and bites during the day so that prevention by bed nets, for example, is ineffective [12].

Obviously, transmission of the disease has been facilitated due to the joint occurrence of large susceptible human populations and competent vectors. However, other aspects are involved in the transmission of arboviruses, since vector abundance and behavior change with the environment. A joint analysis of CHIKV and ZIKV epidemics may provide a better understanding of the commonalities and differences among these two *Aedes*-transmitted diseases. Up to now, these diseases have been studied separately, with a special focus on the reproduction ratio of CHIKV [13, 14, 15, 16] or ZIKV [17, 18, 19, 20]. The uncertainty regarding several parameters, such as the under-reporting ratio and the rate of asymptomatic individuals, have made it difficult to assess the attack rates in naive populations, the relative transmissibility of the viruses, and whether meteorological conditions may alter these parameters.

Here, building on common aspects in location and vectorial transmission, we study in detail the main factors that impacted disease spread. With this objective, we propose a joint model of Chikungunya and Zika transmission based on the time-dependent susceptible-infectious-recovered (TSIR) framework [21], using data from nine distinct territories in French Polynesia and the French West Indies, where both diseases circulated for the first time and caused outbreaks between 2013 and 2016. We also consider the influence of meteorological conditions during the outbreaks. This approach allows to discriminate the respective influence on transmissibility of: (i) properties of each virus itself in a typical situation, (ii) factors that depend on the characteristics of a given area that may be considered as stable over time (e.g. human population density and organization, mobility of humans and vectors, environmental composition), and (iii) weather conditions.

## 2. Material and Methods

We analysed the Zika and Chikungunya epidemics occurring between 2013 and 2016 in six islands or small archipelagoes of French Polynesia and three islands of the French West Indies by fitting weekly incidence data with a common hierarchical transmission model. The two main components of this analysis were: (1) a mechanistic reconstruction of the distribution of the serial interval of the diseases (the time interval between disease onset in a primary and secondary case), including the influence of temperature; and (2) a TSIR model for the generation of observed secondary cases, that included coefficients depending on the territory, the disease and the local weather conditions. The analysis allowed for the identification of the parameters of interest, such as the reporting rate, the reproduction ratio and the attack rate of each outbreak.

### 2.1. Data

Incidence of clinical disease was available from local sentinel networks of health practitioners. Weekly data were collected for the whole duration of the outbreaks, except for the Zika epidemics in the French West Indies that were still ongoing at the time of writing (the period before October 2nd, 2016 was considered for the analysis in that case). In French Polynesia, both Zika and Chikungunya epidemics were monitored by the *Centre d’Hygiène et de Salubrité Publique de Polynésie Française* for six islands and archipelagoes of French Polynesia: the Austral Islands (abbreviated AUS), the Marquesas Islands (MRQ), Mo’orea Island (MOO), the Sous-le-vent Islands (SLV, also named Leeward Islands), Tahiti (TAH), and the Tuamotus (TUA) [22, 23]. In the French West Indies, the two epidemics were monitored by the *CIRE Antilles-Guyane* for Guadeloupe (GUA), Martinique (MRT), and Saint-Martin (STM) [24, 25].

For each disease, we obtained aggregated numbers of suspected cases by week of disease onset for each area. In the case of Saint-Martin, where CHIKV circulated at low intensity for several months after an initial peak, we only considered the first wave of the outbreak (25 weeks). Suspected cases of ZIKV infection were defined as a rash with or without fever and at least two signs among conjunctivitis, arthralgia, or oedema. Suspected cases of CHIKV infection were defined as a fever with arthralgia. As the number of active sentinel centres changed from week to week, we used the projected numbers of cases extrapolated from the number of reporting health practitioners in the area provided by the local authorities [17]. Weather data were collected for each location and outbreak period from the nearest weather station available in the *Weather Underground* website [26]. Daily reports of mean temperature (in °C) and total precipitation (in cm) were aggregated by week on the same grid as incidence data.

### 2.2 Serial interval distribution

We extended the approach used in [13] to obtain the distribution of the serial interval between onset of disease in a primary and secondary case for Zika and Chikungunya. In this framework, the serial interval is split in four parts: (*i*) the infectious period before symptoms in the primary case; (*ii*) the time from infectiousness in the primary case to an infectious mosquito bite; (*iii*) the time from the initial contaminating bite to a transmitting bite in the mosquito – that depends on the extrinsic incubation period (EIP) and the mosquito gonotrophic cycle -; and (*iv*) the incubation period in the secondary case.

The characteristics of each part are well informed in the literature and puts forward the importance of temperature in three respects. First, the duration of the EIP shortens as temperature increases. Following [27, 28], we computed the mean EIP duration at temperature *T* as *κ(T)* = *κ*_28_ × exp(−0.21(*T* − 28)) [29], with base value *κ*_28_ measured at 28°C fixed at 3 days for Chikungunya [30, 31] and at 6 days for Zika [32, 9]. Second, the mean duration of one gonotrophic cycle also decreases with temperature, from about 5 days at 22°C to 2 days at 28°C [33] but increases again at higher temperatures. This association was described using the function γ*(T)* = 56.64 − 3.736 × *T* + 0.064 × *T*^2^ [33]. Third, mosquito mortality rate is affected by temperature, but the effect measured in field studies is very limited within the range of values relevant in this study. Hence, the daily mortality rate was fixed to 0.29 [34].

Taking all information into account, we computed the serial interval distributions as a function of temperature *T*. These were characterized by their mean *μ_SI_(T)*, standard deviation *σ_SI_(T)* and cumulative distribution function *G(t, T)*. Further details are available in the Supplementary Information.

### 2.3 Transmission model

The Bayesian model for analysing outbreaks was based on the model presented by Perkins et al. [21]. Here, we first describe analysis for one disease in one location, then extend the framework to a hierarchical model encompassing CHIKV and ZIKV in all locations.

#### One island, one disease

We are interested in modelling the time series *O = {O_t_}_t=1,…,K_* of the weekly number of incident cases reported to the surveillance systems, where *K* is the duration of the epidemic in weeks. *O* consists of the cases who sought clinical advice and were diagnosed, a fraction of the (unobserved) incident infected cases *I = {I_t_}_t=1,…,K_*. We therefore write, in the "observation" level of the model, that *O_t_* is a proportion *ρ* of all cases *I_t_* according to:

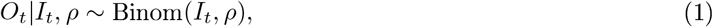

where *ρ* is the probability that an infected case consulted with a health professional, was diagnosed and reported.

In the “transmission” level, we link incidence *I_t_* with past observed incidences *O_t_*^−^ = {*O_1_,…,O_t−1_*} as:

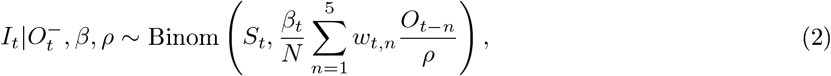

where *N* the total population of the island and *β_t_* is a time-dependent transmission parameter. The term

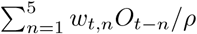

summarizes exposure to infectious mosquitoes at time *t*: it is an average of past incidence with weights defined by the serial interval distribution 
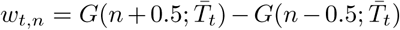
 computed at the mean temperature 
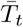
 over the 5 weeks preceding t. The number of susceptible individuals 
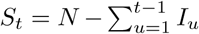

at the beginning of period *t* is computed as 
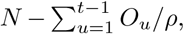
 nothing that *O_u/ρ_* is a first order approximation to *I_u_*.

To avoid data augmentation with the unobserved *I* during estimation, we collapse the "observation" and "transmission" levels into a single binomial distribution:

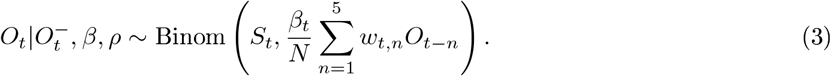

In a final step, we account for the imprecise nature of the *O* data, since observed cases *O* have been extrapolated from limited information provided by a network of local health practitioners. We therefore allow for over-dispersion using a negative binomial distribution instead of the binomial, as:

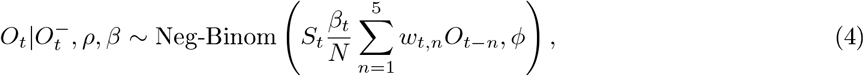

where variance is computed as the mean divided by *ϕ*.

The joint probability of data and parameters is finally

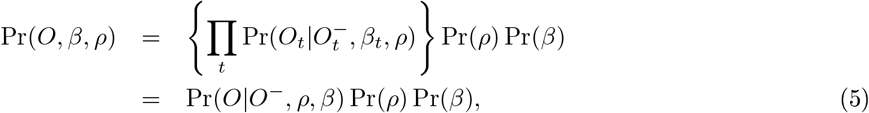

where Pr(*β*) and Pr(*ρ*) are prior distributions described later.

#### Several islands, several diseases

We introduced a hierarchical structure in the model for reporting rates and transmission. Reporting rates *ρ_ij_* in island *i* for disease *j* were modelled using a logistic-normal model

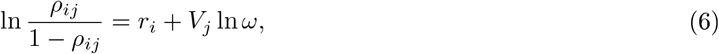

where 
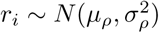

is a random island effect, *V_j_* is 1 for ZIKV and 0 for CHIKV and *ω* is the odds-ratio of reporting ZIKV cases relative to CHIKV cases during an epidemic. Three parameters *θ_ρ_* = {*μ_ρ_*, *σ_ρ_*,*ω*} define the model.

Likewise, we allowed for a random island coefficient in the transmission term and fixed effects for disease and weather covariates, as follows:

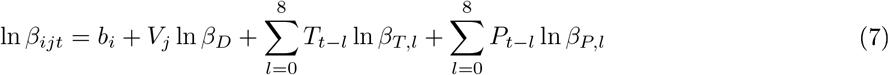

where 
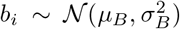

is an island-specific random parameter, *β_D_* is the relative transmission of ZIKV compared to CHIKV, and the last two terms capture the influence of temperature *T* and precipitation *P* for the last nine weeks on transmission with nine transmission effects each, *β_T,l_* and *β_P,l_* - we chose to consider a time lag up to eight weeks based on previous knowledge on dengue ecology [35]. Transmission thus depends on parameters

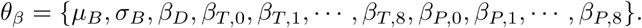

Writing *O_ij_* = {*O_ijt_*} the observed incidence in the outbreak in island *i* and disease *j*, and *O* the whole dataset, we have:

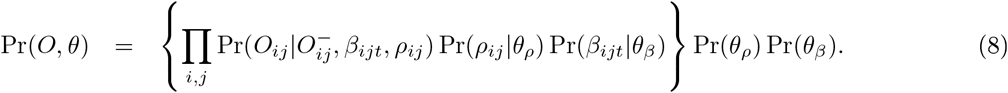

where *θ* = {*θ_ρ_, θ_β_*} is the whole set of parameters to be estimated.

#### Models and outcomes

We defined a set of models to structure our investigation of the effect of disease and weather covariates on transmission:

- the *baseline* model considered a single transmission rate for the entirety of each outbreak irrespective of weather conditions (i.e. parameters *β_T,l_* and *β_P,l_* were set to 1);
- the *free* model, with the same hypotheses as the *baseline* model, but with independent random terms *b_ij_* for each disease instead of the combination *b_i_ + ln β_D_V_j_* in the transmission term, dropping the assumption of dependence between two outbreaks occurring in the same area (see Supplementary Information for more details);
- the *weather* models built on the baseline model by including a dependence of transmission on weather conditions. We considered 3 possibilities: temperature alone (*weather T*), precipitation alone (*weather P*), and both at the same time (*weather T & P*).

Model fits were compared using the leave-one-out information criterion (LOOIC), a Bayesian information criterion especially adapted to hierarchical models [36]. To summarize the intensity of transmission, we computed reproduction ratios based on the general formula 
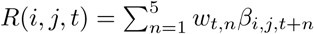
 In particular, we computed for each disease an island-averaged basic reproduction ratio 
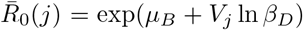
, and for each island a disease-specific basic reproductive ratio *R_0_(i,j)=exp(b_i_+ V_j_ ln β_D_)*.

#### Prior information & estimation

We used thick-tailed, weakly informative prior distributions as shown in Table 1 and in the Supplementary Information[37, 38]. The joint posterior distributions of the parameters were explored with the Hamiltonian Monte Carlo *NUTS* algorithm, as implemented in *Stan* 2.9.0 [39]. We used eight chains with 10,000 iterations each and discarded the first 25% as burn-in. Each chain was sampled every 10-th iteration to reduce autocorrelation. Convergence of the chains was assessed both visually and by inspecting the Gelman-Rubin ratio Ȓ for every estimated parameter. The posterior distributions were summarised by their means and 95% credible intervals (95%CI).

We checked that all parameters were identifiable using simulated datasets (see Supplementary Information). Importantly, we found that even when epidemics were not observed to their end, the results were sensible.

**Table 1:** Prior distributions.

| Parameter | Meaning | Prior distribution |
| --- | --- | --- |
| $\mu_\rho$ | Island-averaged reporting parameter | $\mu_\rho \sim \text{Normal}(\mu = 0, \sigma = 1)$ |
| $\sigma_\rho$ | Between-islands SD in reporting | $\sigma_\rho \sim \text{Inv-Gamma}(\alpha = 10, \beta = 10)$ |
| $\omega$ | Odds-ratio of reporting for ZIKV relative to CHIKV | $\ln \omega \sim \text{Normal}(\mu = 0, \sigma = 1)$ |
| $\mu_B$ | Island-averaged baseline transmission parameter | $\mu_B \sim \text{Student}(\nu = 5, \mu = 0, \sigma = 2.5)$ |
| $\sigma_B$ | Between-islands SD for transmission | $\sigma_B \sim \text{half-Cauchy}(x_0 = 0, \gamma = 2.5)$ |
| $\beta_D$ | Relative transmission of ZIKV with respect to CHIKV | $\ln \beta_D \sim \text{Student}(\nu = 5, \mu = 0, \sigma = 2.5)$ |
| $\beta_{T,l}$ | Relative transmission according to temperature (with time lag $l$) | $\ln \beta_{T,l} \sim \text{Student}(\nu = 5, \mu = 0, \sigma = 2.5)$ |
| $\beta_{P,l}$ | Relative precipitation according to temperature (with time lag $l$) | $\ln \beta_{P,l} \sim \text{Student}(\nu = 5, \mu = 0, \sigma = 2.5)$ |
| $\phi$ | Overdispersion in reported cases | $\phi \sim \text{half-Cauchy}(x_0 = 0, \gamma = 2.5)$ |

## 3. Results

### 3.1 CHIKV and ZIKV outbreaks in French Polynesia and French West Indies

In French Polynesia, ZIKV outbreaks occurred between October, 2013 and March, 2014, followed one year later by CHIKV outbreaks (October, 2014 – March, 2015). Overall, there were about 30,000 observed clinical cases of Zika and 69,000 of Chikungunya, corresponding to an observed cumulated incidence of 11% and 26% (Table 2). The dynamics of the outbreaks were similar in most of the six studied areas, with a steep increase after the first reported cases and a mean outbreak duration of 20 weeks (Fig. 1A). Except in the Austral islands, there were more reports of CHIKV cases than of ZIKV cases (2.1 to 4.5 times more).

**Figure 1:**
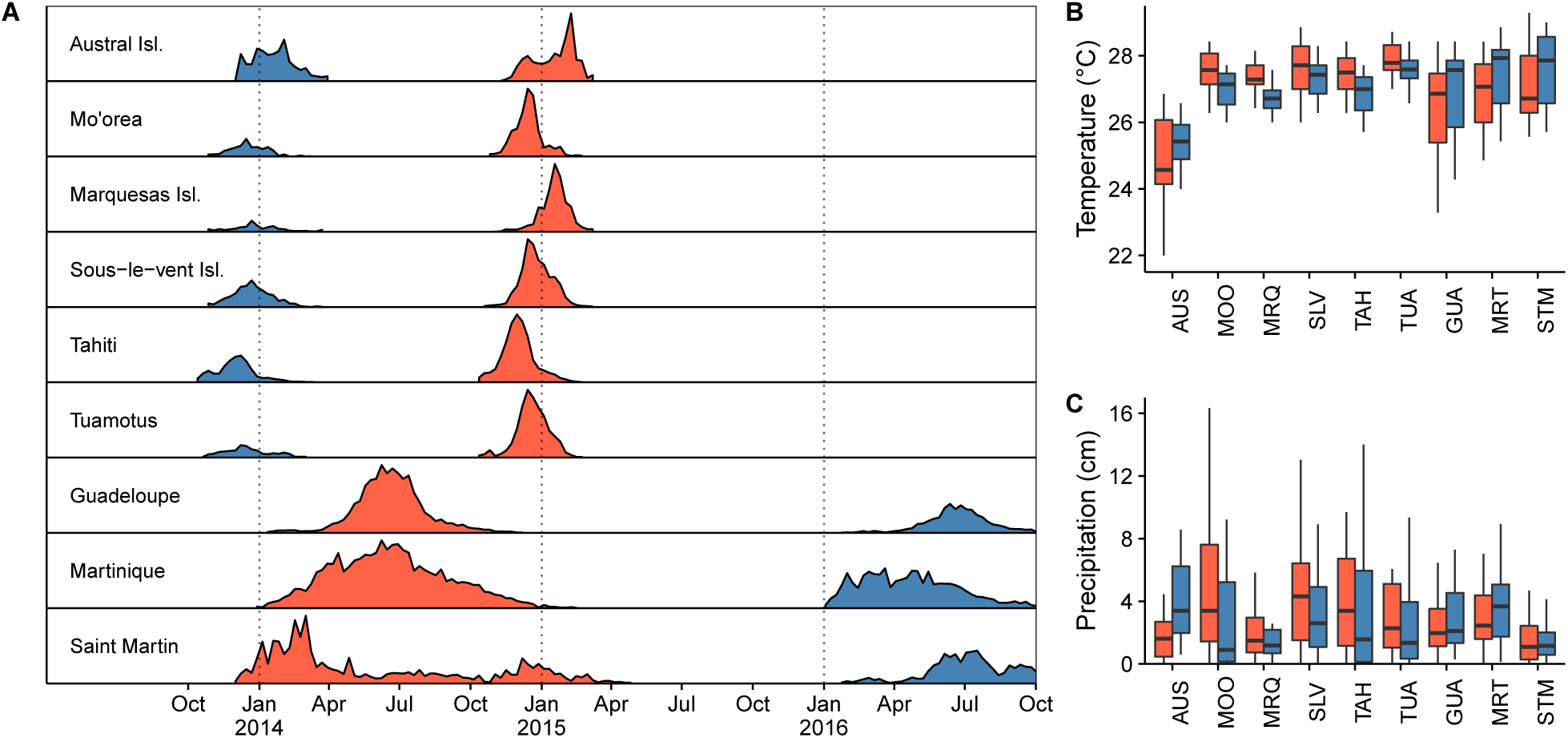
(A) Profiles of CHIKV (red) and ZIKV(blue) outbreaks in the nine territories under study. (B-C) Distributions of weekly mean temperatures (in °C) and precipitation (in cm) during the corresponding epidemic periods.

**Table 2:** Territories included in the 2013-2016 ZIKV and CHIKV outbreaks in French Polynesia and the French West Indies, with the observed cumulated incidence.

| Territory | Territory code | Population | ZIKV cases |  | CHIKV cases |  |
| --- | --- | --- | --- | --- | --- | --- |
|  |  |  | *Number* | *%* | *Number* | *%* |
| *Polynesia* |  |  |  |  |  |  |
| Austral Islands | AUS | 7,000 | 1,208 | 17% | 1,321 | 19% |
| Mo'orea Island | MOO | 16,000 | 1,235 | 8% | 3,830 | 24% |
| Marquesas Islands | MRQ | 9,000 | 994 | 11% | 4,762 | 53% |
| Sous-le-vent Islands | SLV | 33,000 | 3,912 | 12% | 8,046 | 24% |
| Tahiti | TAH | 184,000 | 21,406 | 12% | 45,722 | 25% |
| Tuamotus | TUA | 16,000 | 1,211 | 8% | 5,017 | 31% |
| *West Indies* |  |  |  |  |  |  |
| Guadeloupe | GLP | 400,000 | 30,454<sup>†</sup> | 8% | 81,321 | 20% |
| Martinique | MTQ | 385,000 | 37,295<sup>†</sup> | 10% | 72,539 | 19% |
| Saint-Martin | MAF | 36,000 | 2,519<sup>†</sup> | 7% | 3,309 | 9% |

<sup>†</sup> As of October 2nd, 2016 (still ongoing).

In the French West Indies, CHIKV outbreaks occurred first (between December, 2013 and April, 2015) and ZIKV outbreaks started in January, 2016 and were still ongoing by October, 2016. Overall, there were 159,000 reported clinical cases of Chikungunya and 70,000 of Zika. If we consider only the first epidemic wave in Saint-Martin, the CHIKV epidemics lasted from 25 to 59 weeks, with an observed cumulated incidence between 9% and 20%. As of October 2nd, 2016, i.e. 36 to 39 weeks after the first reported cases, ZIKV was still circulating in the three considered islands at a low pace.

The weather conditions over the outbreak periods showed diverse behavior across islands (Fig. 1B and 1C). The temperature in French Polynesia islands was almost constant around 27-28 °C, except in the Austral

Islands where the temperature was colder (around 24-26 °C). In the West Indies, the range of variation was larger and the average temperature was around 27 °C, slightly higher during the periods of ZIKV circulation. Rainfall showed an opposite trend, with more variation in the Pacific islands than in the West Indies.

### 3.2 Reconstructed temperature-dependent serial interval

The mean serial interval ranged from 1.5 to 2.7 weeks for CHIKV and from 2.2 to 4.7 weeks for ZIKV according to the changes in temperature over the periods (Fig. 2). The changes were limited in most Polynesian islands, reflecting stable temperature over time, except for the Austral islands. On the contrary, the serial interval tended to be longer on average and more variable in the West Indies.

**Figure 2:**
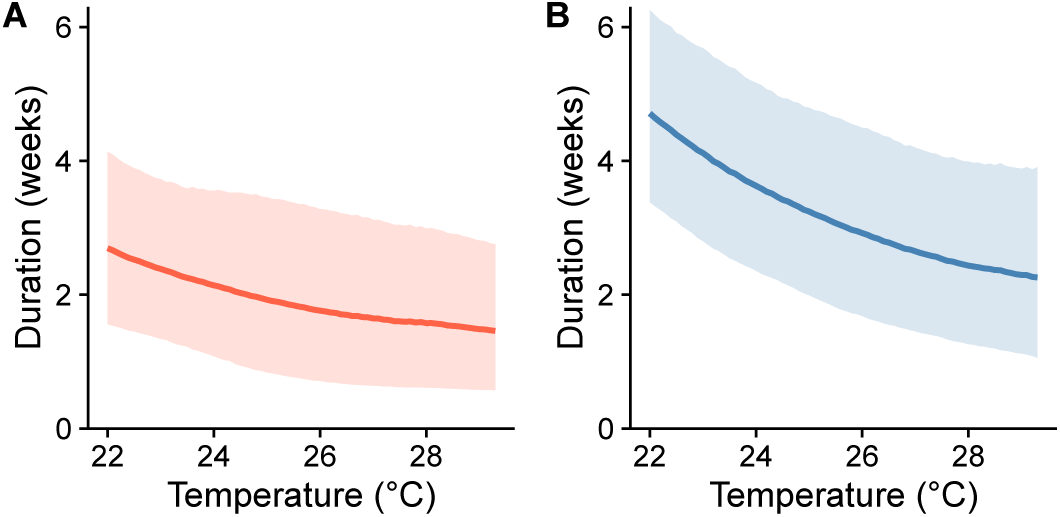
Mean, 2.5% and 97.5% quantiles of the serial interval duration for CHIKV (A) and ZIKV (B) according to temperature (in °C).

### 3.3 Model fit and parameter estimates

The *baseline* model captured the essential characteristics of the outbreaks, as shown in Fig. 3. Overall, the timecourse of the CHIKV outbreaks was fitted more accurately than that of the ZIKV outbreaks: some predicted peaks in incidence were slightly off target, for example in Tahiti or the Marquesas Islands. The t was also very good for the three ZIKV epidemics still under way in the French West Indies. The *baseline* model, with an additive effect of disease and island on transmission performed similarly to the *free* model according to the LOOIC (difference = +5; standard error = 9), suggesting that this description of transmission was indeed adequate. The two models yielded similar *R*_0_ values, despite dierences for some Zika epidemics and in general larger fluctuations predicted by the *free* model (see Supplementary Information).

**Figure 3:**
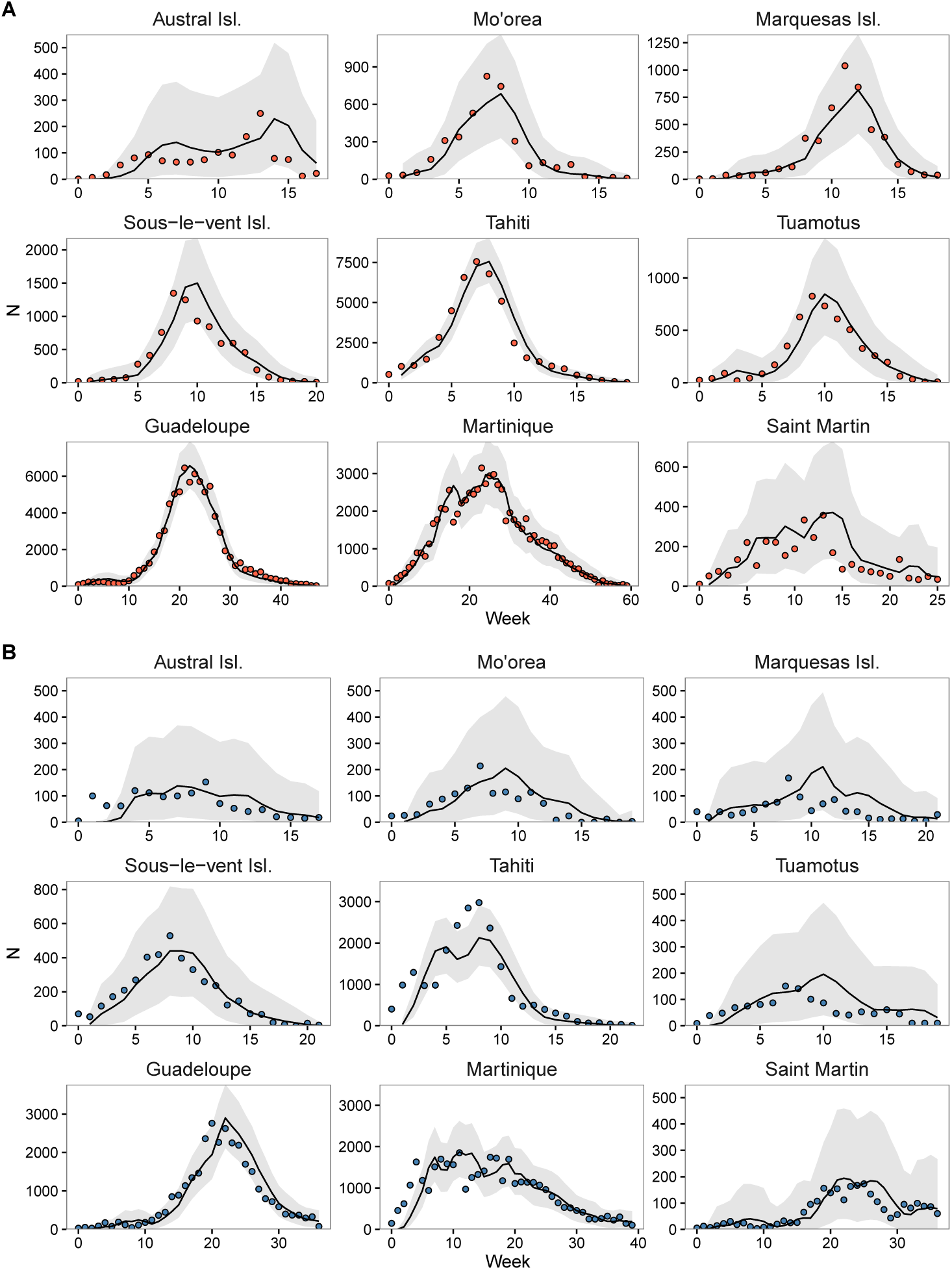
Observed numbers of clinical cases (circles) for CHIKV (panel A) and ZIKV (panel B), and corresponding one-week ahead predicted values for each week obtained from the baseline model using the observed numbers in the preceding weeks and the posterior distributions of the parameters (the lines correspond to mean values; the grey zones correspond to 95% credible intervals).

The parameters of the *baseline* model are reported in Table 3. The island-averaged reporting rate 
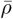
 differed according to the disease, estimated at 40% [29%; 54%] for CHIKV and 19% [12%; 28%] for ZIKV. The transmissibility ratio between ZIKV and CHIKV was _D_ = 1.04 [95%CI: 0.97; 1.13], showing no significant difference in transmissibility between the diseases. Indeed, the island-averaged reproductive ratio was 
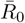
 = 1.80 [1.54; 2.12] for CHIKV and 1.88 [1.59; 2.22] for ZIKV.

**Table 3:**
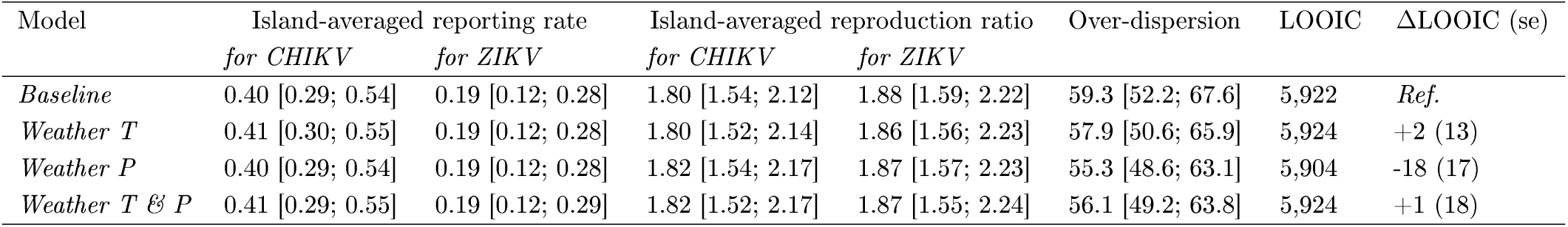
Posterior means and 95% credible intervals for the parameters of the four models.

The variance for the random island effects in reporting and in transmission were different from 0, indicating heterogeneity between locations (Fig. 4). The island-specific reproduction ratios *R*_0_ were lower in the French West Indies compared to French Polynesia, especially in Martinique. The island-specific reporting rates grouped islands in two sets, the first with high reporting rates, around 50% for CHIKV and 25% for ZIKV, and including the Austral Islands, the Marquesas Islands, the Tuamotus, and Martinique, and a second group with smaller reporting rates, around 25% for CHIKV and 15% for ZIKV, including Guadeloupe, Saint-Martin, and the other Polynesian islands.

**Figure 4:**
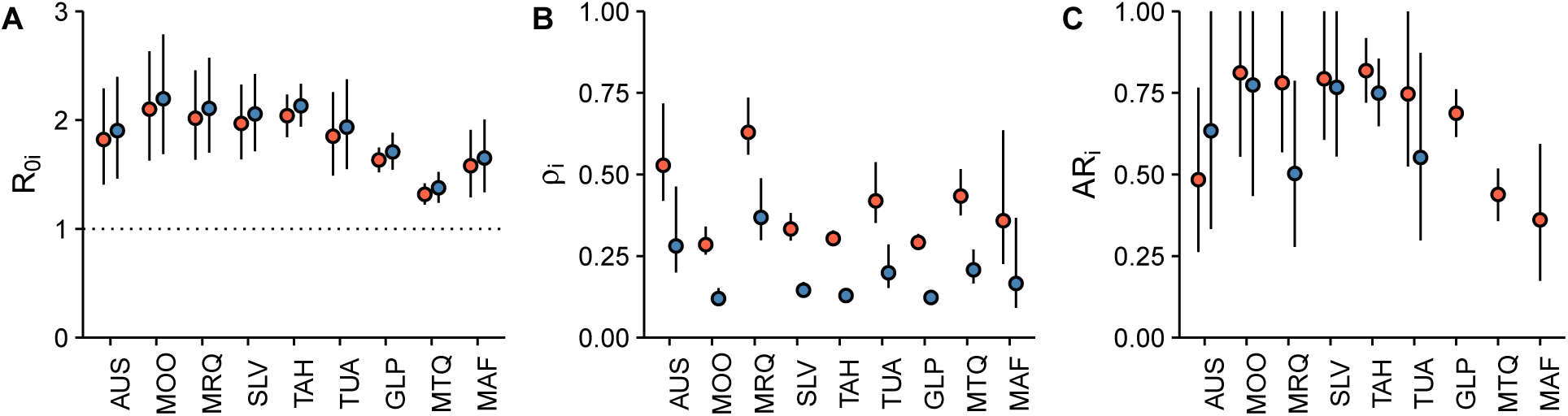
(A) Island-specic basic reproduction ratio *R*_0*i*_; (B) reporting rate *ρ*_*i*_; and (C) final attack rate *AR_i_* for CHIKV (red) and ZIKV (blue) and in nine French territories.

Estimated values for the final attack rates follow the spatial pattern of *R*_0_ estimates, exception made for the Zika epidemics in Marquesas Islands and the Tuamotus, for which we found substantially smaller values. Attack rates for both diseases were as high as 70-80% in Mo'orea, the Sous-le-vent Islands, Tahiti, and Guadeloupe, while closer to 50% for the Austral Islands, Martinique and Saint Martin. With the exception of the Austral Islands, Zika caused smaller outbreaks.

### 3.4 Weather conditions and disease transmission

We added all combinations of precipitation and temperature on transmission and compared models using LOOIC values. Including precipitation in transmission (model *weather P*) improved the t (LOOIC difference = -18, standard error = 17), while the effect of temperature was not meaningful (Table 3). In particular, including temperature alone in the transmission term suggested a small reduction in transmission with higher temperature four weeks in the past, but this effect disappeared when precipitation and temperature were entered at the same time in the model (Fig 5A). On the contrary, the effect of local precipitation showed a marked pattern on transmission, with higher rainfall decreasing transmission one to two weeks later and then increasing transmission four to six weeks afterwards (Fig 5B), with or without adjustment on temperature. For a typical increase in average weekly precipitation of 1 cm, this corresponded to a reduction of transmission by 10% in the short term and an increase of about 20 to 40% four to six weeks afterwards. We repeated the analyses including one lag at a time, instead of all nine together, and found essentially the same pattern, indicating no problems due to autocorrelation.

**Figure 5.**
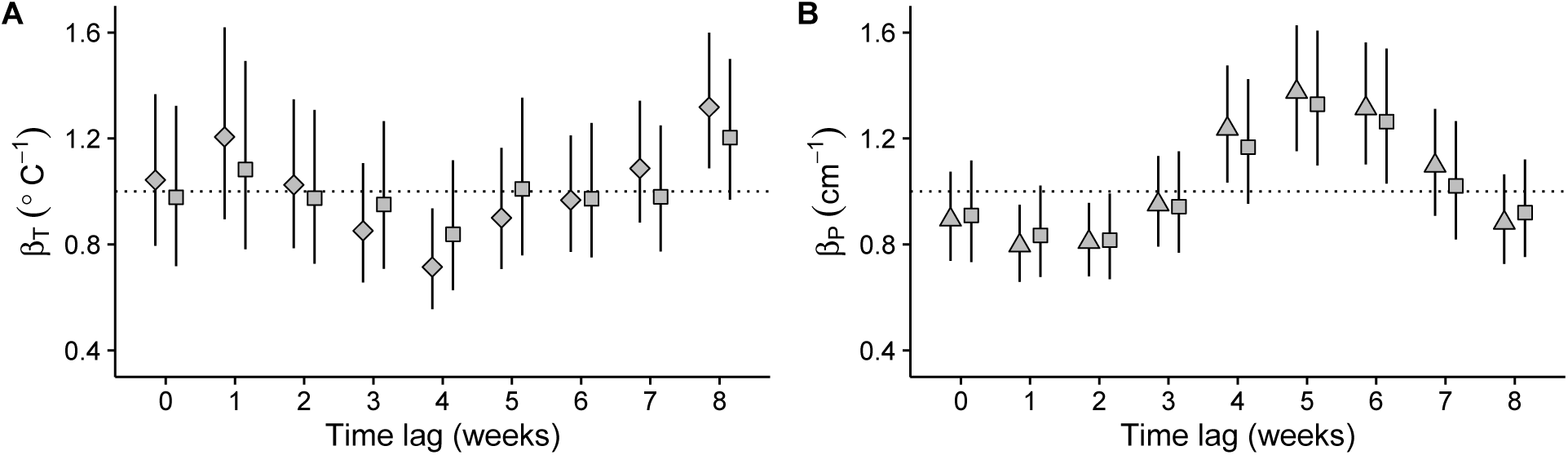
Effect of a variation of 1°C of the weekly-averaged mean temperature (panel A) and of 1 cm of the weekly-averaged precipitation (panel B) on transmissibility according to a given time lag for the three *weather* models.

Using model *weather P*, the ZIKV epidemic in Tahiti was better explained, with precipitation leading to a better description of the epidemic peak, (Fig. 6), and we also noticed a steepest reduction in the Sous-le-vent CHIKV Islands after the peak. In other islands, the fit was little affected by transmissibility change with weather. Other parameters were not affected by inclusion of weather: reporting rates in the *weather* models were estimated at the same values as in the *baseline* model.

**Figure 6.**
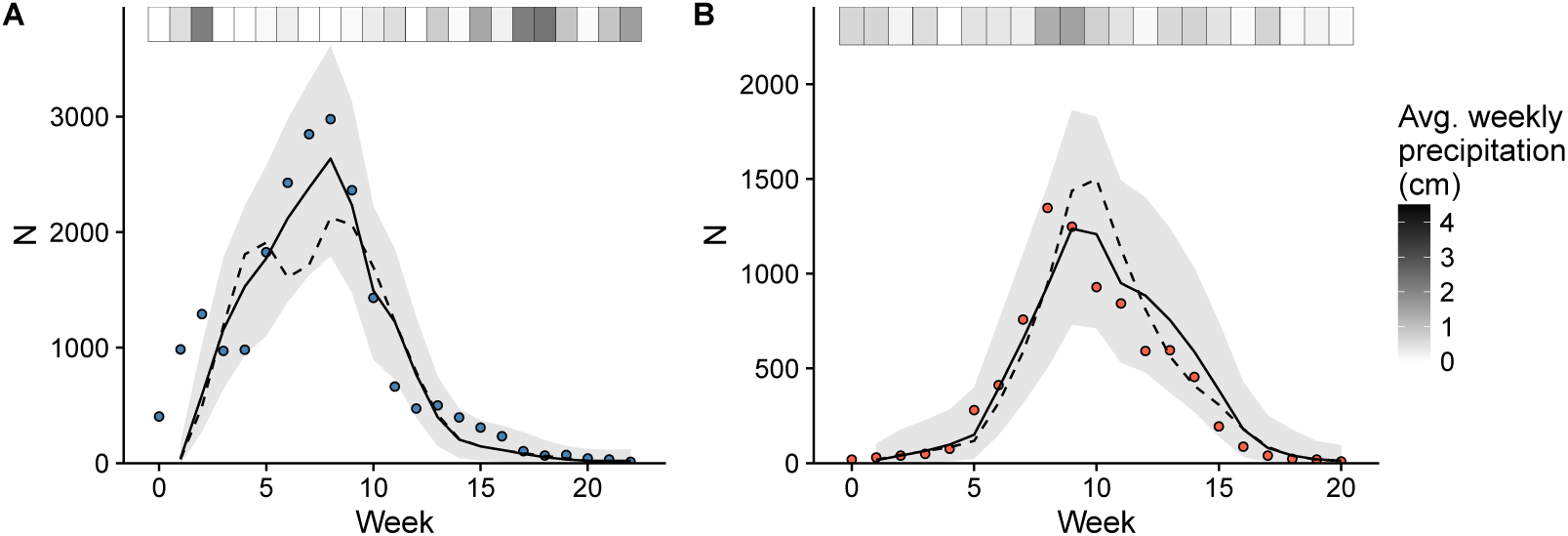
Comparison of the fits of the *baseline* model (dashed line) and of the *weather* model including precipitation (solid line) for the ZIKV outbreak in Tahiti (panel A) and for the CHIKV outbreak in the Sous-le-vent Islands (panel B). The top band shows weekly precipitation.

## 4. Discussion

The rapid spread of arboviral diseases has raised public health concern worldwide. Understanding the drivers of transmission and the sources of variability in the different patterns of propagation requires investigating the effects of viruses and environmental variables, but is complicated by differences in surveillance systems. The hierarchical model here developed combined epidemiological data on the Chikungunya and Zika outbreaks in nine different territories, making optimal use of the available information and allowing for disentangling the role of different components on transmission, thus providing a better understanding of the arboviral dynamics.

Transmission was adequately described with additive contributions of island and virus effects, compared to the less structured *free* model. Then, once the effects of the locality and the weather conditions were taken into account, the difference in transmission between the two viruses was minute. This suggests that the epidemic dynamics are controlled less by the difference between viruses than by factors related to the locality such as mosquito abundance, the local environmental, socio-economic and meteorological conditions. Similar transmissibility is also compatible with reproduction ratio estimates reported for both diseases [17, 19, 18, 40, 41, 21], although we acknowledge that differences in hypotheses (e.g. serial intervals) make such comparisons uneasy. Only one study compared directly the transmissibility of two arboviruses focused on Dengue and Zika epidemics in Yap Island [42], reporting higher transmissibility for Zika.

Besides providing a comparison between viruses, we also assessed variation in transmission with location. Overall, transmission was lower in the French West Indies than in the Pacific islands, although with substantial geographical heterogeneity. Previous analyses reported variation between territories [17, 19, 18, 40]. Kucharski et al. [17] analysed the ZIKV epidemic in French Polynesia with results similar to those reported here: Tahiti, Sous-le-vent and Mo'orea had larger reproduction ratios than Australes and Tuamotu. In the French West Indies, the reproduction ratios previously estimated by [40] for CHIKV suggested higher transmission in Guadeloupe than in Martinique, as reported here. Interestingly, in the West Indies, similar patterns were noticed for CHIKV and ZIKV: in Martinique both outbreaks started quickly after introduction, while in Guadeloupe we observed a delay of a few weeks following introduction, after which the growth was more intense. Patterns of human mobility and commuting [43], housing and lifestyle [44], land use [45], water management and waste collection [46] may explain such differences, and should be further assessed.

Second, we quantified the role of precipitation and temperature in driving transmission for the two diseases, by modelling their effect on transmissibility. We found that precipitation, but not temperature, influenced the epidemics in French Polynesia and French West Indies. Indeed, the difference in LOOIC between the *weather P* model and the *baseline* model was larger than typical chance variation as measured by the standard error. This shows the better predictive accuracy of a model involving the effect of precipitation, although the baseline model already captured the essential features of the epidemics. Including a temperature-dependent serial interval distribution was not responsible for the lack of association with temperature, as the same results were found with a temperature independent serial interval distribution (see Supplementary Information). It is possible that temperature effects within the range spanned in the study (quite narrow and close to the transmissibility optimum [47, 21]) are too small to be detected by our statistical approach. The effect of increased rainfall was twofold, reducing transmission two weeks later but enhancing it with a time lag of four to six weeks. These effects were the same when the lagged terms were introduced one at a time rather than simultaneously in the model, and persisted irrespective of the inclusion of temperature. The effect of weather on arboviral diseases has been the subject of extensive research, mainly on Dengue fever [48, 49, 35, 50, 51, 52, 53], and, with a focus on climatic rather than local meteorological factors, on Chikungunya [21]. These studies yielded mixed results, although most reported a positive association between precipitation and incidence, explained by an increased abundance of vectors [49, 35, 51, 52, 53]. Natural rainfall can certainly create puddles allowing oviposition and *Ae. ageypti*'s larval development, and our results are in line with this interpretation, the five weeks lag matching the time required for the maturation of larvae and the incubation periods [35]. This may not be the case in all places, since artificial water storage reduces the impact of such natural phenomena [54, 35]. In the present analysis we found that the inclusion of precipitation improved the t of the epidemics of Zika in Tahiti and Chikungunya in Sous-le-vent Islands, while other outbreaks were less affected by the inclusion of this ingredient, as precipitation was more stable during the epidemic period. The short term negative association between increased rainfall and transmission is less commonly reported [48, 55, 56, 57, 58]. It may be due to a reduced exposure to mosquitoes either because adult mosquitos are more likely to die [59] or because the human population would reduce outdoor activity and be protected by indoor mechanisms (e.g. air-cooling) [54].

A third finding was the dissimilarity in the reporting ratios between the two diseases and between islands. We estimated that overall only 19% of all Zika cases ended up being reported, against 40% of Chikungunya cases. These estimates are consistent with the much higher asymptomatic rate of ZIKV infections (around 80%) compared to CHIKV infections (around 30%) reported in serological studies in Yap Island and French Polynesia [60, 61]. Differences between territories may be ascribed to the organization of the local surveillance systems in relation with the structure of the target population, where a major difficulty is the assessment of the actual population covered [62]. The operational characteristics of surveillance networks in these islands are unknown, however increased medical participation is often possible in the less populated islands and may contribute to larger reporting rates estimated in the smaller territories of Austral Islands, Marquesas Islands, the Tuamotus, and Saint Martin (Fig. 4).

The values for the final attack rates show the same spatial structure as the reproduction numbers for CHIKV. For ZIKV, however, *R*_0_ was not always a good predictor of the final attack rate of the epidemic, as was noted elsewhere [60]. For instance, Marquesas Islands and the Tuamotus show higher values of *R*_0_ than the Austral Islands and, at the same time, smaller attack rates. Data on the Zika outbreaks were noisy in these territories, thus the use of the information from the Chikungunya outbreaks in the same locations showed here its strongest effect, reducing the indetermination and guiding the estimation of the epidemic parameters. In the French West Indies, the model was able to capture the distinctive dynamics at play in Guadeloupe and Martinique, two very similar islands that displayed different epidemic profiles. We predicted that, despite similar numbers of CHIKV cases were reported in the two territories, the difference in the outbreak behavior was associated with a higher attack rate in Guadeloupe than Martinique (66% vs. 45%), reflecting the delayed but more intense transmission in the former island. Zika is showing a similar pattern suggesting that stable properties of the environment, the population and its spatial distribution may be responsible for the observed behavior. Further research conducted at a smaller scale would be needed to better understand the causes of this difference. Differences in transmission and reporting between islands were described here using a random effect model. Using additional information, it would be possible to actually model these differences as a function of island characteristics. Local characteristics such as economic indicators, land usage, commuting, could be considered in a further investigation.

Our model presented some innovative features. Its hierarchical structure allowed for the epidemic assessment in several territories and for two diseases at the same time while keeping low the number of parameters (e.g. random island factors are drawn from a common distribution). By relying on other outbreaks occurring in the same area and on outbreaks in other areas, we obtained stable, precise posterior estimates for the reproductive ratios, the reporting rates and the attack rates of each epidemic. In particular, joint estimation of transmissibility and reporting rate within a territory was possible because we hypothesized no pre-existing immunity to the diseases in the populations and fixed the serial interval. Transmissibility was therefore determined by initial exponential growth, and reporting rate by the difference between actual and model-predicted attack rates. The whole model was also tested with simulated data, and was able to precisely recover the initial values (see Supplementary Information). Some limitations, however, need to be acknowledged. It was not possible to estimate the serial interval distributions alongside basic reproductive ratio and reporting rate. We overcame this issue by developing a mechanistic model of the serial interval based on combined information from different bibliographic sources. Still, uncertainty remained on some parameters, particularly the association between the EIP and temperature, which was based on Dengue experiments [29], following the approach of [27, 28]. It is important to notice, however, that a sensitivity analysis ignoring this association yielded very similar results. Another limitation of this analysis is the neglect of sexual transmission, which has been reported in the case of Zika. We argue that, when a competent vector is abundant as in the territories under study, sexual transmission is unlikely to contribute significantly to transmission. Indeed, other works estimated that it accounted for only 3% of transmission [63, 41]. Finally, we made the assumption that *Ae. aegypti* is the main vector involved in transmission, and calibrated the distribution of the serial interval accordingly.

In conclusion, we jointly analysed epidemics of Zika and Chikungunya in nine island territories to quantify the respective roles of the virus, the territory and the weather conditions in the outbreak dynamics. We showed that Chikungunya and Zika have similar transmissibility when spreading in the same location. Accounting for the level of precipitation improved the modelling of the epidemic profiles, notably for the outbreaks of Zika in Tahiti and of Chikungunya in the Sous-le-vent Islands. Eventually, different probabilities of developing symptoms for the two diseases translated in substantial differences in reporting rates. The present study provides valuable information for the assessment and projection of *Aedes*-borne infections spread, by quantifying the impact of virus, territory and weather in transmission. In addition, it introduces an approach that can be adopted in other comparative analyses involving multiple arboviruses and locations.

## Acknowledgements

We thank both the *CIRE Antilles-Guyane* and the *Centre d’Hygiène et de Salubrité Publique de Polynésie Française* for collecting the data and making it publicly available.

### Author contributions

JR, CP and PYB designed research, interpreted results and wrote the manuscript. JR performed the data analysis.

### Declaration of interest

We declare that we have no conflict of interest.

## References

[1] WHO, Zika Virus Outbreak Global Response Interim Report (May 2016). URL http://apps.who.int/iris/bitstream/10665/207474/1/WHO_ZIKV_SRF_16.2_eng.pdf?ua=1.

[2] D. J. Gubler, The changing epidemiology of yellow fever and dengue, 1900 to 2003: full circle?, Comparative immunology, microbiology and infectious diseases 27 (5) (2004) 319–330.

[3] P. Renault, J.-L. Solet, D. Sissoko, E. Balleydier, S. Larrieu, L. Filleul, C. Lassalle, J. Thiria, E. Rachou, de Valk, et al., A major epidemic of chikungunya virus infection on reunion island, france, 2005–2006, The American journal of tropical medicine and hygiene 77 (4) (2007) 727–731.

[4] M. R. Duffy, T.-H. Chen, W. T. Hancock, A. M. Powers, J. L. Kool, R. S. Lanciotti, M. Pretrick, M. Marfel, S. Holzbauer, C. Dubray, L. Guillaumot, A. Griggs, M. Bel, A. J. Lambert, J. Laven, O. Kosoy, A. Panella, B. J. Biggerstaff, M. Fischer, E. B. Hayes, Zika Virus Outbreak on Yap Island, Federated States of Micronesia, New England Journal of Medicine 360 (24) (2009) 2536–2543. doi:10.1056/NEJMoa0805715.URL http://dx.doi.org/10.1056/NEJMoa0805715.

[5] S. C. Weaver, M. Lecuit, Chikungunya Virus and the Global Spread of a Mosquito-Borne Disease, New England Journal of Medicine 372 (13) (2015) 1231–1239. doi:10.1056/NEJMra1406035. URL http://dx.doi.org/10.1056/NEJMra1406035.

[6] D. Musso, D. J. Gubler, et al., Zika virus: following the path of dengue and chikungunya?, The Lancet 386 (9990) (2015) 243–244.

[7] Q. Zhang, K. Sun, M. Chinazzi, A. Pastore-Piontti, N. E. Dean, D. P. Rojas, S. Merler, D. Mistry, Poletti, L. Rossi, M. Bray, M. E. Halloran, I. M. Longini, A. Vespignani, Projected spread of Zika virus in the Americas, bioRxiv (2016) 066456 doi:10.1101/066456. URL http://biorxiv.org/content/early/2016/07/29/066456.

[8] V. Richard, T. Paoaafaite, V.-M. Cao-Lormeau, Vector competence of aedes aegypti and aedes polynesiensis populations from french polynesia for chikungunya virus, PLOS Negl Trop Dis 10 (5) (2016) e0004694.

[9] M. I. Li, P. S. J. Wong, L. C. Ng, C. H. Tan, Oral susceptibility of singapore aedes (stegomyia) aegypti (linnaeus) to zika virus, PLoS Negl Trop Dis 6 (8) (2012) e1792.

[10] J. E. Brown, C. S. McBride, P. Johnson, S. Ritchie, C. Paupy, H. Bossin, J. Lutomiah, I. Fernandez-Salas, A. Ponlawat, A. J. Cornel, et al., Worldwide patterns of genetic differentiation imply multiple ‘domestications’ of aedes aegypti, a major vector of human diseases, Proceedings of the Royal Society of London B: Biological Sciences 278 (1717) (2011) 2446–2454.

[11] E. P. Lima, M. H. S. Paiva, A. P. de Araújo, E. Da Silva, U. M. da Silva, L. N. de Oliveira, A. Santana, C. N. Barbosa, C. C. de Paiva Neto, M. Goulart, et al., Insecticide resistance in aedes aegypti populations from ceará, brazil, Parasit Vectors 4 (5) (2011) 2–12.

[12] S. Christophers, et al., Aedes aegypti (l.) the yellow fever mosquito: its life history, bionomics and structure., Rickard.

[13] P.-Y. Boëlle, G. Thomas, E. Vergu, P. Renault, A.-J. Valleron, A. Flahault, Investigating Transmission in a Two-Wave Epidemic of Chikungunya Fever, Réunion Island, Vector-Borne and Zoonotic Diseases 8 (2) (2008) 207–218. doi:10.1089/vbz.2006.0620. URL http://online.liebertpub.com/doi/abs/10.1089/vbz.2006.0620.

[14] P. Poletti, G. Messeri, M. Ajelli, R. Vallorani, C. Rizzo, S. Merler, Transmission Potential of Chikungunya Virus and Control Measures: The Case of Italy, PLoS ONE 6 (5) (2011) e18860. doi:10.1371/journal.pone.0018860. URL http://dx.doi.org/10.1371/journal.pone.0018860.

[15] L. Yakob, A. C. A. Clements, A Mathematical Model of Chikungunya Dynamics and Control: The Major Epidemic on Réunion Island, PLoS ONE 8 (3) (2013) e57448. doi:10.1371/journal.pone.0057448. URL http://dx.doi.org/10.1371/journal.pone.0057448.

[16] M. Robinson, A. Conan, V. Duong, S. Ly, C. Ngan, P. Buchy, A. Tarantola, X. Rodó, A model for a chikungunya outbreak in a rural Cambodian setting: implications for disease control in uninfected areas, PLoS Negl Trop Dis 8 (9) (2014) e3120. doi:10.1371/journal.pntd.0003120.

[17] A. J. Kucharski, S. Funk, R. M. Eggo, H.-P. Mallet, W. J. Edmunds, E. J. Nilles, Transmission dynamics of zika virus in island populations: a modelling analysis of the 2013–14 french polynesia outbreak, PLOS Negl Trop Dis 10 (5) (2016) e0004726.

[18] C. Champagne, D. G. Salthouse, R. Paul, V.-M. Cao-Lormeau, B. Roche, B. Cazelles, Structure in the variability of the basic reproductive number (r0) for zika epidemics in the pacific islands, eLife 5 (e19874) doi:10.7554/eLife.19874. URL https://elifesciences.org/content/5/e19874.

[19] H. Nishiura, R. Kinoshita, K. Mizumoto, Y. Yasuda, K. Nah, Transmission potential of zika virus infection in the south pacic, International Journal of Infectious Diseases 45 (2016) 95–97. doi:http://dx.doi.org/10.1016/j.ijid.2016.02.017. URL http://www.sciencedirect.com/science/article/pii/S1201971216000370.

[20] G. Chowell, D. Hincapie-Palacio, J. Ospina, B. Pell, A. Tariq, S. Dahal, S. Moghadas, A. Smirnova, L. Simonsen, C. Viboud, Using Phenomenological Models to Characterize Transmissibility and Forecast Patterns and Final Burden of Zika Epidemics, PLoS Currents doi:10.1371/currents.outbreaks.f14b2217c902f453d9320a43a35b9583. URL http://currents.plos.org/outbreaks/?p=68353.

[21] T. A. Perkins, C. J. E. Metcalf, B. T. Grenfell, A. J. Tatem, Estimating Drivers of Autochthonous Transmission of Chikungunya Virus in its Invasion of the Americas, PLoS Curr 7. doi:10.1371/currents.outbreaks.a4c7b6ac10e0420b1788c9767946d1fc.

[22] Directionde la santé, Bureau de veille sanitaire, Surveillance et veille sanitaire en Polynésie Française (Mar. 2015). URL http://www.hygiene-publique.gov.pf/IMG/pdf/bss_sem_9-10-2015-version_mars_2015.pdf.

[23] Centre d’hygiène et de salubrité publique de Polynésie française, Surveillance de la dengue et du zika en Polynésie française (Mar. 2014). URL http://www.hygiene-publique.gov.pf/IMG/pdf/bulletin_dengue_28-03-14.pdf.

[24] CIRE Antilles Guyane, Le point épidémiologique n ° 2 (3 2015). URL http://www.invs.sante.fr/fr/content/download/104810/376847/version/85/file/pe_chikungunya_antilles_060315.pdf.

[25] CIRE Antilles Guyane, Le point épidémiologique n ° 38 (10 2016). URL http://invs.santepubliquefrance.fr/fr/content/download/130573/466695/version/133/file/pe_zika_antilles_guyane_061016.pdf.

[26] Weather underground website (2016). URL https://www.wunderground.com/.

[27] M. A. Johansson, A. M. Powers, N. Pesik, N. J. Cohen, J. E. Staples, Nowcasting the spread of chikungunya virus in the americas, PloS one 9 (8) (2014) e104915.

[28] T. Alex Perkins, A. S. Siraj, C. W. Ruktanonchai, M. U. G. Kraemer, A.J. Tatem, Model-based projections of Zika virus infections in childbearing women in the Americas, Nature Microbiology 1 (2016) 16126. doi:10.1038/nmicrobiol.2016.126. URL http://www.nature.com/articles/nmicrobiol2016126.

[29] M. Chan, M. A. Johansson, The incubation periods of dengue viruses, PloS one 7 (11) (2012) e50972.

[30] M. Dubrulle, L. Mousson, S. Moutailler, M. Vazeille, A.-B. Failloux, Chikungunya virus and aedes mosquitoes: saliva is infectious as soon as two days after oral infection, PloS one 4 (6) (2009) e5895.

[31] M. Dupont-Rouzeyrol, V. Caro, L. Guillaumot, M. Vazeille, E. D'Ortenzio, J.-M. Thiberge, N. Baroux, A.-C. Gourinat, M. Grandadam, A.-B. Failloux, Chikungunya virus and the mosquito vector aedes aegypti in new caledonia (south pacific region), Vector-Borne and Zoonotic Diseases 12 (12) (2012) 1036–1041.

[32] J. Boorman, J. Portereld, A simple technique for infection of mosquitoes with viruses transmission of zika virus, Transactions of the Royal Society of Tropical Medicine and Hygiene 50 (3) (1956) 238–242.

[33] L. B. Carrington, M. V. Armijos, L. Lambrechts, C. M. Barker, T. W. Scott, Effects of fluctuating daily temperatures at critical thermal extremes on aedes aegypti life-history traits, PLoS One 8 (3) (2013) e58824.

[34] O. J. Brady, M. A. Johansson, C. A. Guerra, S. Bhatt, N. Golding, D. M. Pigott, H. Delatte, M. G. Grech, P. T. Leisnham, R. Maciel-de Freitas, et al., Modelling adult aedes aegypti and aedes albopictus survival at different temperatures in laboratory and field settings, Parasites & vectors 6 (1) (2013) 1–12.

[35] R. Barrera, M. Amador, A. J. MacKay, Population Dynamics of Aedes aegypti and Dengue as Influenced by Weather and Human Behavior in San Juan, Puerto Rico, PLOS Negl Trop Dis 5 (12) (2011) e1378. doi:10.1371/journal.pntd.0001378. URL http://journals.plos.org/plosntds/article?id=10.1371/journal.pntd.0001378.

[36] A. Vehtari, A. Gelman, J. Gabry, Practical bayesian model evaluation using leave-one-out cross-validation and waic, arXiv preprint arXiv:1507.04544.

[37] A. Gelman, J. B. Carlin, H. S. Stern, D. B. Rubin, Bayesian data analysis, Vol. 2, Chapman & Hall/CRC Boca Raton, FL, USA, 2014.

[38] A. Gelman, et al., Prior distributions for variance parameters in hierarchical models (comment on article by browne and draper), Bayesian analysis 1 (3) (2006) 515–534.

[39] B. Carpenter, A. Gelman, M. Hoffman, D. Lee, B. Goodrich, M. Betancourt, M. A. Brubaker, J. Guo, P. Li, A. Riddell, Stan: a probabilistic programming language, Journal of Statistical Software.

[40] S. Cauchemez, M. Ledrans, C. Poletto, P. Quenel, H. De Valk, V. Colizza, P. Boëlle, Local and regional spread of chikungunya fever in the americas, Euro surveillance: bulletin Europeen sur les maladies transmissibles= European communicable disease bulletin 19 (28) (2014) 20854.

[41] D. Gao, Y. Lou, D. He, T. C. Porco, Y. Kuang, G. Chowell, S. Ruan, Prevention and control of zika as a mosquito-borne and sexually transmitted disease: A mathematical modeling analysis, Scientific Reports 6 (2016) 28070.

[42] S. Funk, A. J. Kucharski, A. Camacho, R. M. Eggo, L. Yakob, W. J. Edmunds, Comparative analysis of dengue and zika outbreaks reveals differences by setting and virus, PLoS Negl Trop Dis 10 (2) (2016) e0005173. doi:10.1371/journal.pntd.0005173.

[43] S. T. Stoddard, B. M. Forshey, A. C. Morrison, V. A. Paz-Soldan, G. M. Vazquez-Prokopec, H. Astete, R. C. Reiner, S. Vilcarromero, J. P. Elder, E. S. Halsey, et al., House-to-house human movement drives dengue virus transmission, Proceedings of the National Academy of Sciences 110 (3) (2013) 994–999.

[44] P. Reiter, S. Lathrop, M. Bunning, B. Biggerstaff, D. Singer, T. Tiwari, L. Baber, M. Amador, J. Thirion, J. Hayes, C. Seca, J. Mendez, B. Ramirez, J. Robinson, J. Rawlings, V. Vorndam, S. Waterman, D. Gubler, G. Clark, E. Hayes, Texas Lifestyle Limits Transmission of Dengue Virus, Emerging Infectious Diseases 9 (1) (2003) 86–89. doi:10.3201/eid0901.020220. URL http://wwwnc.cdc.gov/eid/article/9/1/02-0220_article.htm.

[45] E. F. Lambin, A. Tran, S. O. Vanwambeke, C. Linard, V. Soti, Pathogenic landscapes: interactions between land, people, disease vectors, and their animal hosts, International journal of health geographics 9 (1) (2010) 1.

[46] T. P. Monath, Dengue: the risk to developed and developing countries, Proceedings of the National Academy of Sciences 91 (7) (1994) 2395–2400.

[47] E. Mordecai, J. Cohen, M. V. Evans, P. Gudapati, L. R. Johnson, K. Miazgowicz, C. C. Murdock, J. R. Rohr, S. J. Ryan, V. Savage, M. Shocket, A. S. Ibarra, M. B. Thomas, D. P. Weikel, Temperature determines Zika, dengue and chikungunya transmission potential in the Americas, bioRxiv (2016) 063735 doi:10.1101/063735. URL http://biorxiv.org/content/early/2016/07/15/063735.

[48] S. Banu, W. Hu, C. Hurst, S. Tong, Dengue transmission in the Asia-Pacific region: impact of climate change and socio-environmental factors, Tropical Medicine & International Health 16 (5) (2011) 598–607. doi:10.1111/j.1365-3156.2011.02734.x. URL http://onlinelibrary.wiley.com/doi/10.1111/j.1365-3156.2011.02734.x/abstract.

[49] C. F. Li, T. W. Lim, L. L. Han, R. Fang, Rainfall, abundance of Aedes aegypti and dengue infection in Selangor, Malaysia, The Southeast Asian Journal of Tropical Medicine and Public Health 16 (4) (1985) 560–568.

[50] B. C. Ho, K. L. Chan, Y. C. Chan, Aedes aegypti (L.) and Aedes albopictus (Skuse) in Singapore City, Bulletin of the World Health Organization 44 (5) (1971) 635–641. URL http://www.ncbi.nlm.nih.gov/pmc/articles/PMC2427847/.

[51] G. L. Sia Su, Correlation of Climatic Factors and Dengue Incidence in Metro Manila, Philippines, AMBIO: A Journal of the Human Environment 37 (4) (2008) 292–294. doi:10.1579/0044-7447(2008) 37[292:COCFAD]2.0.CO;2. URL http://www.bioone.org/doi/abs/10.1579/0044-7447(2008)37%5B292:COCFAD%5D2.0.CO%3B2.

[52] L. Lu, H. Lin, L. Tian, W. Yang, J. Sun, Q. Liu, Time series analysis of dengue fever and weather in Guangzhou, China, BMC Public Health 9 (2009) 395. doi:10.1186/1471-2458-9-395. URL http://dx.doi.org/10.1186/1471-2458-9-395.

[53] V. Wiwanitkit, An observation on correlation between rainfall and the prevalence of clinical cases of dengue in Thailand, Journal of Vector Borne Diseases 43 (2) (2006) 73–76.

[54] C. C. Jansen, N. W. Beebe, The dengue vector Aedes aegypti: what comes next, Microbes and Infection 12 (4) (2010) 272–279. doi:10.1016/j.micinf.2009.12.011. URL http://www.sciencedirect.com/science/article/pii/S1286457910000109.

[55] Y. L. Hii, H. Zhu, N. Ng, L. C. Ng, J. Rocklöv, Forecast of Dengue Incidence Using Temperature and Rainfall, PLoS Negl Trop Dis 6 (11) (2012) e1908. doi:10.1371/journal.pntd.0001908. URL http://dx.doi.org/10.1371/journal.pntd.0001908.

[56] P.-C. Wu, J.-G. Lay, H.-R. Guo, C.-Y. Lin, S.-C. Lung, H.-J. Su, Higher temperature and urbanization affect the spatial patterns of dengue fever transmission in subtropical Taiwan, The Science of the Total Environment 407 (7) (2009) 2224–2233. doi:10.1016/j.scitotenv.2008.11.034.

[57] H. Halide, P. Ridd, A predictive model for Dengue Hemorrhagic Fever epidemics, International Journal of Environmental Health Research 18 (4) (2008) 253–265. doi:10.1080/09603120801966043.

[58] S. Thammapalo, V. Chongsuwiwatwong, D. McNeil, A. Geater, The climatic factors influencing the occurrence of dengue hemorrhagic fever in Thailand, The Southeast Asian Journal of Tropical Medicine and Public Health 36 (1) (2005) 191–196.

[59] F. Fouque, R. Carinci, P. Gaborit, J. Issaly, D. J. Bicout, P. Sabatier, Aedes aegypti survival and dengue transmission patterns in French Guiana, Journal of Vector Ecology 31 (2) (2006) 390–399. doi:10.3376/1081-1710(2006)31[390:AASADT]2.0.CO;2. URL http://www.bioone.org/doi/abs/10.3376/1081-1710%282006%2931%5B390%3AAASADT%5D2.0.CO%3B2.

[60] J. Lessler, L. H. Chaisson, L. M. Kucirka, Q. Bi, K. Grantz, H. Salje, A. C. Carcelen, C. T. Ott, J. S. Sheffield, N. M. Ferguson, D. A. T. Cummings, C. J. E. Metcalf, I. Rodriguez-Barraquer, Assessing the global threat from Zika virus, Science (2016) aaf8160 doi:10.1126/science.aaf8160. URL http://science.sciencemag.org/content/early/2016/07/13/science.aaf8160.

[61] J. E. Staples, R. F. Breiman, A. M. Powers, Chikungunya fever: an epidemiological review of a re-emerging infectious disease, Clinical Infectious Diseases 49 (6) (2009) 942–948.

[62] C. Souty, C. Turbelin, T. Blanchon, T. Hanslik, Y. Le Strat, P.-Y. Boëlle, Improving disease incidence estimates in primary care surveillance systems, Population health metrics 12 (1) (2014) 1.

[63] C. L. Althaus, N. Low, How Relevant Is Sexual Transmission of Zika Virus?, PLOS Medicine 13 (10)(2016) e1002157. doi:10.1371/journal.pmed.1002157. URL http://journals.plos.org/plosmedicine/article?id=10.1371/journal.pmed.1002157.

